# Oxidative stress alters transcript localization of disease-causing genes in the retinal pigment epithelium

**DOI:** 10.1101/2021.01.07.425741

**Authors:** Tadeusz J. Kaczynski, Elizabeth D. Au, Michael H. Farkas

**Author notes:** **Correspondence:** Michael Farkas.

## Abstract

Nuclear retention is a mechanism whereby RNA transcripts are held in the nucleus to maintain a proper nuclear-to-cytoplasmic balance or as a stockpile for use in responding to stimuli. Many mechanisms are employed to determine whether transcripts are retained or exported to the cytoplasm, though the extent to which tissue- or cell-type, stressors, or disease pathogenesis affect this process remains unclear. As the most biochemically active tissue in the body, the retina must mitigate endogenous and exogenous stressors to maintain cell health and tissue function. Oxidative stress, believed to contribute to the pathogenesis, or progression, of age-related macular degeneration (AMD) and inherited retinal dystrophies (IRDs), is produced both internally from biochemical processes, as well as externally from environmental insult. To evaluate the effect of oxidative stress on transcript localization in the retinal pigment epithelium (RPE), we performed poly-A RNA sequencing on nuclear and cytoplasmic fractions from induced pluripotent stem cell-derived retinal pigment epithelium (iPSC-RPE) cells exposed to hydrogen peroxide, as well as untreated controls. Under normal conditions, the number of mRNA transcripts retained in the nucleus exceeded that found in studies of other tissues. Further, the nuclear-to-cytoplasmic ratio of transcripts is altered following oxidative stress, as is the retention of genes associated with AMD, IRDs, and those important for RPE physiology. These results provide a retention catalog of all expressed mRNA in iPSC-RPE under normal conditions and after exposure to hydrogen peroxide, offering insight into one of the potential roles oxidative stress plays in the progression of visual disorders.

## Introduction

It has long been widely accepted that messenger RNA (mRNA) is transcribed, spliced, processed, and rapidly exported from the nucleus to the cytoplasm. Given the role of mRNA to produce a protein, it seems vital that export to the cytoplasm rapidly occur to promote translation. This view, however, is being challenged by studies of ever-increasing sophistication (Bahar Halpern et al., 2015). On a cell-by-cell basis, transcription occurs in bursts. For any given gene, one cell may express an excess of the transcript compared to the neighboring cell. Additionally, to maintain a homeostatic environment, excess transcripts can be retained in the nucleus, some for a majority of their lifetime (Bahar Halpern et al., 2015). Nuclear RNA retention, on a global scale, is a relatively new concept. It is not yet completely clear how transcripts are retained, if retention differs by cell- or tissue-type, and how stress or disease affect this process.

While nuclear retention studies are still in their infancy, some have begun to examine the role played by a variety of factors. Aberrant RNA localization can be caused by internal factors such as mutations in nuclear export factors, triplet repeat expansion, and mis-splicing leading to intron retention (Wegener and Müller-McNicoll, 2018). These factors are implicated in a multitude of neurodegenerative diseases, such as various types of myotonic dystrophy, Huntington’s disease, and glial cell tumors (Comincini et al., 2006; Larkin and Fardaei, 2001; Mastroyiannopoulos et al., 2010; Sun et al., 2015). Repeat expansions in *DMPK* and *ZNF9* that are causes of muscular dystrophy lead to the aberrant mRNA being retained in nuclear foci (Mastroyiannopoulos et al., 2012). Here, the mutant mRNAs recruit RNA binding proteins, thus preventing their normal function in the cell.

External factors have also been shown to play a role in nuclear retention. Export of mRNA under normal circumstances is performed with strict quality control, but heat stress causes changes in mRNA export from the nucleus (Zander et al., 2016). Typically, an mRNA is checked for proper splicing, capping and polyadenylation prior to being exported to the cytoplasm. However, in times of heat stress, heat shock responsive mRNAs bypass quality control checks and are rapidly exported to the cytoplasm for translation. On the other hand, export of mRNAs not required to respond to the stressor is inhibited. Other external stressors remain to be studied; specifically of note with regard to this study, little is known about the role of oxidative stress on mRNA export.

Though studies have indeed begun to uncover a potential role for nuclear retention in disease pathogenesis, more analysis is necessary. Retinal disease is especially complex, due to genetic heterogeneity and the lack of complete understanding of both the underlying genetic components and environmental effects. Age-related macular degeneration (AMD) results in the loss of the retinal pigment epithelium (RPE) in the macula region of the eye, leading to death of the overlying photoreceptors and central vision loss. While no specific genetic mutations have been identified to cause AMD, 34 genetic loci have been implicated with increased risk via genome-wide association studies (Fritsche et al., 2016). The genetic component of AMD accounts for only 37% of disease pathology, while environmental components (oxidative stress, smoking, and diet) account for the remainder. Moreover, inherited retinal degenerations (IRDs) are caused by mutations in over 250 genes that lead to loss of either photoreceptors or RPE and eventual blindness (Daiger et al., 2013). The mutations alone can lead to vision loss, but it is believed that oxidative stress plays a role in exacerbating the phenotype. The retina is a highly metabolic tissue, producing intrinsic reactive oxygen, but is also subject to external stimuli (light, cigarette smoke, diet) that increases the load of reactive oxygen. In the case of IRDs, the cells are already compromised due to the underlying disease-causing mutations and this may lead to an inability to cope with the reactive oxygen. Oxidative stress is known to cause a wide array of challenges to autophagy, endoplasmic reticulum function, protein folding, and mitochondrial function (Bhat et al., 2015; Cao and Kaufman, 2014; Cui et al., 2012; Filomeni et al., 2015; Kiffin et al., 2006; Malhotra and Kaufman, 2007). The effect of oxidative stress on RNA retention/export of known retinal disease-causing genes and genes important for defining the RPE and maintaining proper cell function (herein referred to as RPE markers) has not been studied.

By studying the whole coding transcriptome, with particular emphasis on genes important to maintaining RPE function and those known to contribute to retinal disease, we can determine the role oxidative stress plays on their retention or export from the nucleus. Using RNA-Seq, we show that oxidative stress globally affects RNA localization, and this is especially evident in both the RPE markers and retinal disease-causing genes.

## Materials and Methods

### 1. Maintenance and differentiation of iPSCs

All reagents were purchased from Invitrogen (Carlsbad, CA) unless noted otherwise. Human iPSC (line ATCC-BXS0114, ATCC, ACS-1028) were seeded at 500,000 cells in a 10-cm dish coated with Matrigel (Corning). Cells were maintained in TeSR-E8 media (Stem Cell Technologies) with Rock Inhibitor (Y-27632 dihydrochloride, Santa Cruz Biotechnology) at a final concentration of 1 μM/mL. Media without Rock Inhibitor was changed daily. The procedure for differentiating human iPSCs toward RPE was performed as previously described (Au et al., 2017; Gamm and Meyer, 2010). The culture was maintained in RDM until day 80, when RPE was dissected and passaged. This process was performed 5 times to generate the 5 technical replicates used for this study.

### 2. Cell fractionation

Subcellular fractionation was carried out as in Rio et al. 2010 with minor adjustments (Rio et al., 2010). Briefly, iPSC-RPE cells were incubated either in RDM media (untreated samples) or in media with 500 μM hydrogen peroxide (treated samples) for 3 hours, then washed three times with phosphate buffered saline (PBS), incubated at 37°C for 5 minutes with Tryple Express dissociation reagent (Fisher Scientific, Cat#: 12-605-010), and collected via scraping. Cells were pelleted via centrifugation and resuspended in ice-cold cell disruption buffer (10 mM KCl, 1.5 mM MgCl2, 20 mM Tris-HCl [pH 7.5], 1 mM dithiothreitol [DTT, added just before use]). To facilitate swelling, cells were incubated on ice for 20 minutes, then transferred to an RNase-free dounce homogenizer. Homogenization was achieved using 15-20 strokes of the pestle, and the homogenate was visualized under a microscope to ensure that greater than 90% of the cell membranes were sheared while the nuclei remained intact. Residual cytoplasmic material was separated from the nuclei by adding 0.1% Triton X-100 and mixing gently by inversion. The nuclei were pelleted via centrifugation, and the supernatant (containing the cytoplasmic fraction) transferred to a new tube. The nuclear pellet was washed in 1 mL of ice-cold cell disruption buffer, and both the nuclear and cytoplasmic fractions were centrifuged. The cytoplasmic supernatant was transferred to a new tube and the wash was removed and discarded from the nuclear pellet.

### 3. RNA isolation

RNA was isolated from the nuclear and cytoplasmic fractions using Tri-Reagent (Molecular Research Center Inc., Cat#: TR 118) following the manufacturer’s protocol. The nuclear and cytoplasmic samples were mixed well by inversion, transferred to phase-lock heavy tubes (VWR, Cat#: 10847-802), and incubated at room temperature for 5 minutes. Chloroform (200 μL) was added to each sample, followed by vigorous mixing for 15 seconds and then 15 minute incubation at room temperature. Samples were centrifuged, and the aqueous (top) phase transferred to a new 1.5 mL microcentrifuge tube. To remove any contaminating phenol, 400 μL chloroform was added, vigorously mixed, incubated at room temperature for 2 minutes, and centrifuged. The aqueous phase, containing the RNA, was transferred to a new 1.5 mL microcentrifuge tube. Each volume of RNA solution was then thoroughly mixed with one 10^th^ volume of 3M sodium acetate, 1 volume of isopropanol, and 2.5 μL RNA-grade glycogen. Samples were incubated at −80°C for 1 hour to precipitate RNA, centrifuged to pellet the RNA, and the supernatant was discarded. To wash the RNA pellets, 75% ethanol was added, then briefly vortexed and centrifuged. Ethanol was removed and this wash was repeated. Following the second wash, ethanol was removed and pellets were to air dried for 5-10 minutes. RNA was resuspended in 22 μL DNase-free, RNase-free water and quantified via nanodrop spectrophotometer.

### 4. RNA library preparation and sequencing

RNA-sequencing libraries were prepared using the SureSelect Strand-Specific RNA Library Prep for Illumina Multiplexed Sequencing kit (Agilent, Cat#: G9691A) according to the manufacturer’s protocol using 100 ng total RNA, and each sample was indexed for multiplexing. Prior to sequencing, library quality and quantity were determined using High Sensitivity Screen Tape on a TapeStation 4150 (Agilent). Sequencing was performed using a NextSeq 500 (Illumina) generating 2 x 75 bp reads.

### 5. Sequencing analysis

Reads were aligned to the human genome (build hg38) with STAR v2.5.2b, and counts generated for all transcripts in gencode v29 using Rsubread v1.32.4 (Dobin et al., 2013; Harrow et al., 2012; Liao et al., 2019). Counts were normalized using reads per kilobase per million reads (rpkm) to examine overall expression. Differential expression analysis using DESeq was performed to determine localization and effect of treatment (Anders and Huber, 2012).

### 6. RNA-fluorescent in situ hybridization

ARPE-19 cells were seeded onto coverslips in a 48-well plate and grown to confluence. Coverslips were prepared using the ViewRNA Cell Plus Assay Kit (Fisher Scientific, Cat#: 88-19000-99) according to the manufacturer’s protocol with the minor alteration of fixation and permeabilization using 3:1 methanol:glacial acetic acid at room temperature. To stain the nuclei, the coverslips were incubated in Hoechst solution. The cells were then mounted on slides for visualization using a Leica TCS SPE confocal microscope.

## Results

### 1. Whole coding transcriptome localization and validation

To examine RNA localization under normal conditions, we generated human induced pluripotent stem cell derived retinal pigmented epithelium (iPSC-RPE) cells using the BXS0114 iPS cell line. Nuclear and cytoplasmic RNA was isolated from five technical replicates of each, and RNA-Seq libraries were prepared. Each sample was sequenced on an Illumina NextSeq 500, and generated an average of 20 million reads, with 86% of reads uniquely aligned to the hg38 human genome build, and, 92% of aligned reads uniquely aligning to one genomic location.

Overall gene expression analysis revealed that approximately one-third of all transcripts are expressed (RPKM > 0.5) in each sample type, with significant overlap between nuclear and cytoplasmic fractions. In total, 76,249 coding transcripts are expressed, 71,086 of these are expressed in the nuclear fraction, and 66,309 in the cytoplasmic fraction. Additionally, to confirm that the iPSC-RPE are sufficiently RPE-like, we examined expression of a comprehensive list of RPE markers, originally developed by Liao et al. (2010) (Liao et al., 2010). Of the 86 RPE markers, 83 were expressed in the iPSC-RPE (FIGURE S1).

To examine localization, we compared transcript expression in the two fractions, and considered a transcript to be localized if it had at least 2-fold greater expression in one fraction, with an adjusted p-value < 0.01. If a transcript had a fold-change less than 1.9 in either direction, it was considered to be mixed localization, regardless of p-value. Any transcript not meeting these criteria was not assigned a specific localization status. Our analysis identified 3,048 coding transcripts localized to the nucleus, 2,129 to the cytoplasm, and 43,431 mixed. (Figure 1A).

**Figure 1.**
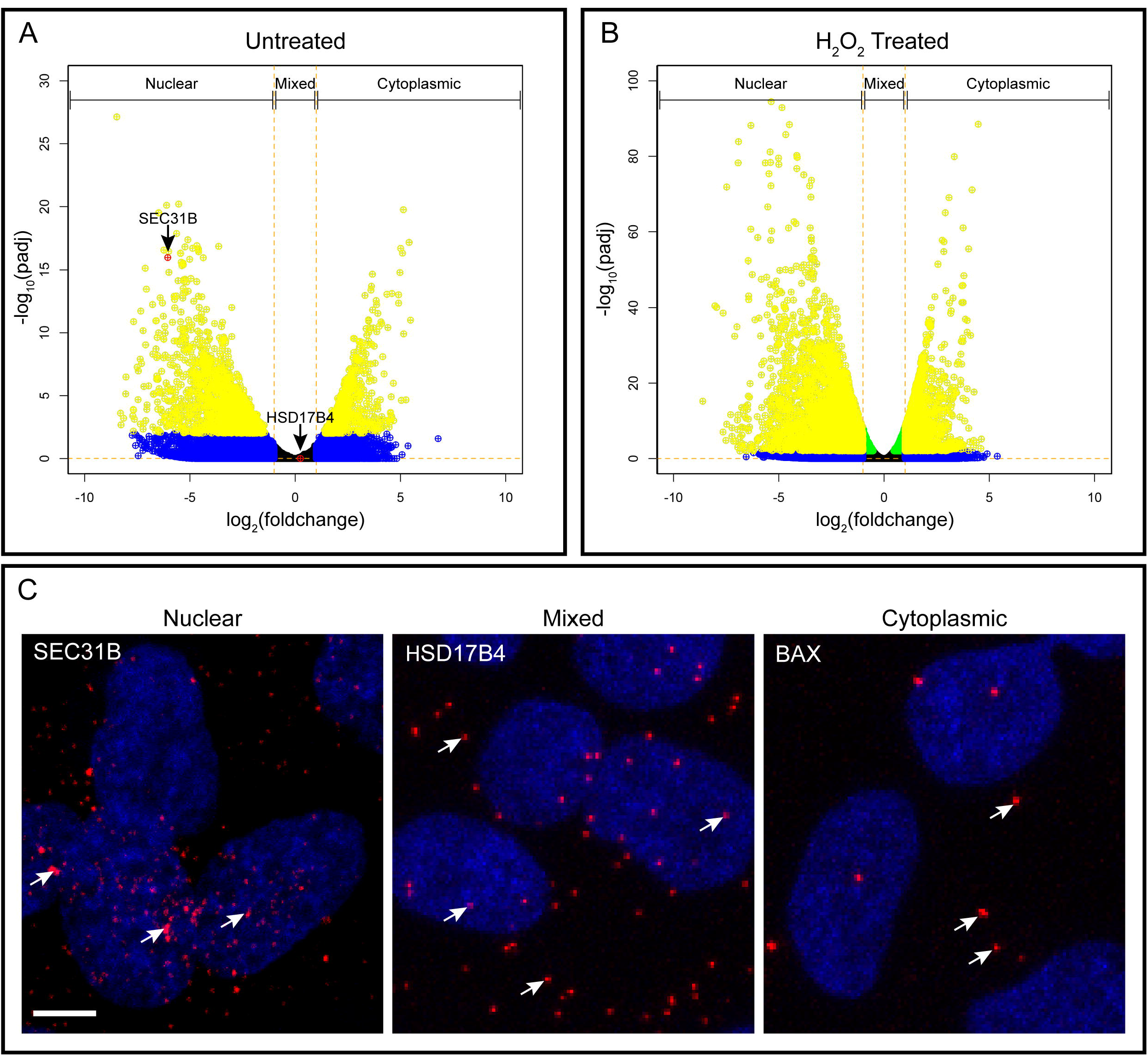
Distribution of coding transcripts in the RPE. Volcano plots of all transcripts in control BXS iPSC-RPE (**A**) and H2O2-treated BXS iPSC-RPE (**B**). Log_2_ cytoplasm:nuclear fold change and corresponding log10 adjusted p-value are plotted for each transcript. Transcripts with fold change >2 are colored blue, adjusted p<0.01 are green, both fold change >2 and adjusted p<0.01 are yellow. Genes confirmed via FISH are red (**A**). Note that BAX is not expressed in the nuclear fraction, hence has infinite fold change and is not seen on the plot. (**C**) RNA-FISH images confirming localization of SEC31B, BAX, and HSD17B4 (red) and counterstained with Hoechst solution (blue). Arrows indicate some of the localized RNAs. Scale bar is 5 μm.

RNA fluorescent in situ hybridization (RNA-FISH) is the gold-standard for determining RNA localization (Alexander and Devaraj, 2017; Beckman et al., 2018; Cui et al., 2016). Accordingly, we performed RNA-FISH using probes targeted to transcripts that were determined by RNA-Seq to have nuclear (SEC31B, ENST00000492667.5, cyto:nuc expression ratio = 0.015), cytoplasmic (BAX, ENST00000515540.5, cyto:nuc expression ratio = Inf), and mixed (HSD17B4, ENST00000256216.11, cyto:nuc expression ratio = 1.2) localization (Figure 1C). The RNA-FISH assays were conducted in ARPE-19 cell line, which is frequently used to model native RPE. In all cases, the RNA-FISH results validated the RNA-Seq findings.

### 2. Effect of oxidative stress on localization and expression

To determine the influence of oxidative stress on RNA localization, we performed the same experiments as above following treatment of the iPSC-RPE with 500 μM H2O2 for 3 hours. Each sample was sequenced on an Illumina NextSeq 500, and generated an average of 14 million reads, with 86% of reads uniquely aligned to the hg38 human genome build, and, 92% of aligned reads uniquely aligning to one genomic location. We again generated depth of coverage plots, and showed that this was sufficient to fully cover the transcriptome.

Overall expression analysis in the treated samples revealed similar, though slightly lower numbers compared to the control. In total, 73,669 transcripts were expressed in the H2O2-treated line, with 69,294 expressed in the nuclear fraction, and 63,581 expressed in the cytoplasmic fraction. Further, we again detected 83 of the 86 RPE markers to be expressed (Figure S1).

While overall expression does not significantly change after treatment, we found H2O2 caused a transcriptome-wide increase in localization. Over twice as many transcripts were localized to the nucleus and cytoplasm in the treated line (7,995 and 4,813, respectively), though the number of mixed localization transcripts (44,462) remained similar to that seen in control (Figure 1B). Notably, 2,376 of the nuclear localized, and 1,286 of the cytoplasmic localized transcripts are shared between both control and treated (Figure 2). In other words, the majority of transcripts localized to one fraction in the control remain so after peroxide treatment, with 78% of the nuclear and 60% of the cytoplasmic transcripts maintaining their localization after treatment. Rather than seeing a shift in localization, we are seeing an influx of additional localized transcripts after treatment.

**Figure 2.**
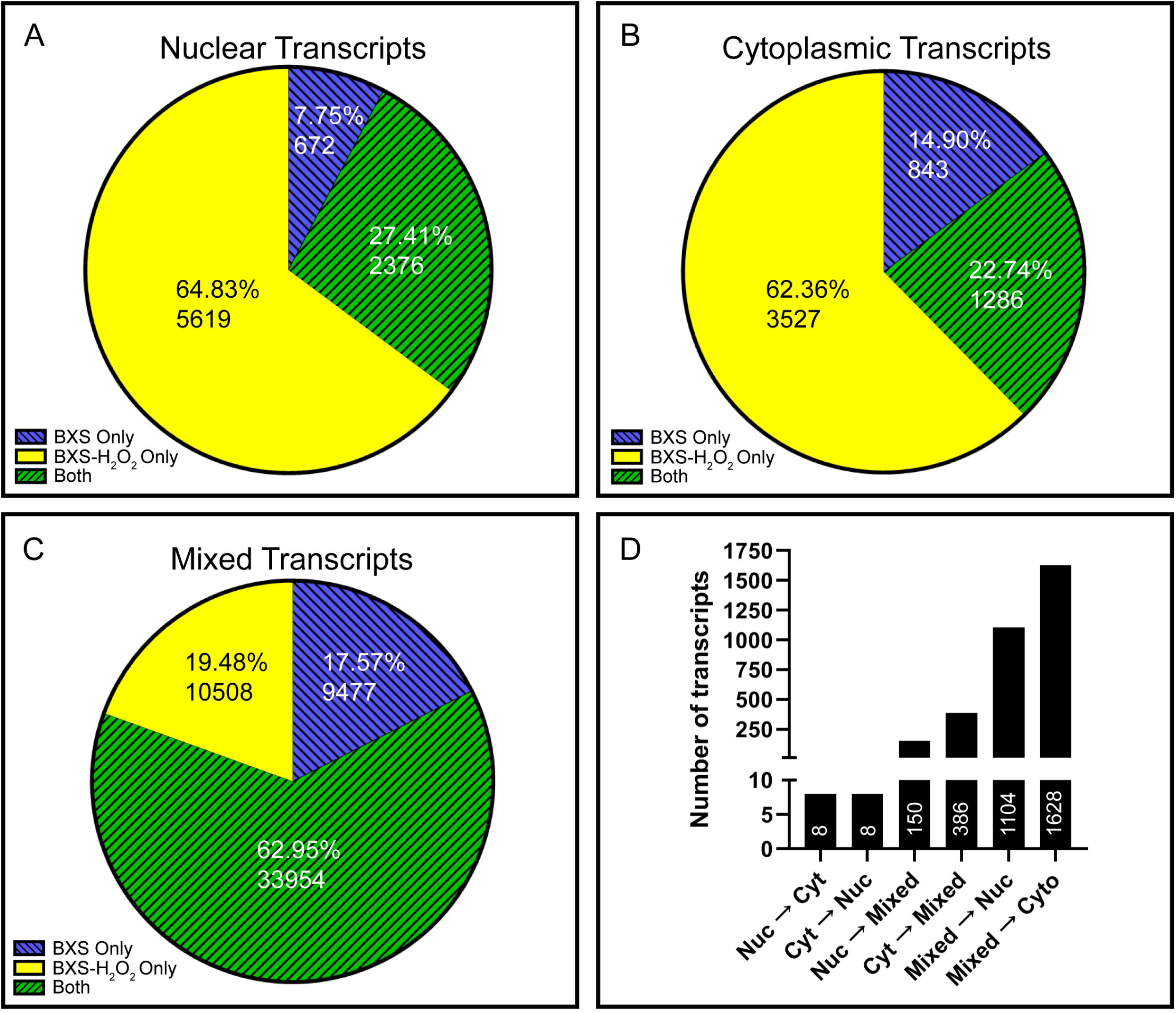
RPE transcript localization is altered by oxidative stress. All transcripts localized to the nucleus (**A**), cytoplasm (**B**), or showing mixed localization (**C**) in control samples (blue), treated samples (yellow), or both (green). Bar graph shows the number of transcripts with altered localization after treatment (**D**). Transitions from control to treated are indicated (e.g. Nuc → Cyt indicates transcripts that are nuclear localized in the control samples and cytoplasmically localized in the treated samples).

To better understand the overall effect on localization resulting from H2O2 treatment, we examined the changes in greater detail. In only a few cases does the localization change from cytoplasmic to nuclear, or vice versa, after treatment with H2O2. Indeed, only 16 transcripts are localized in one fraction under control conditions and change to the other fraction after H2O2, with 8 moving in each direction. Slightly more transcripts are localized to one fraction in control samples and classified as mixed after H2O2 treatment. We find 150 transcripts to be nuclear localized in the control cells and mixed after treatment, and 386 localized to the cytoplasm in control and mixed after treatment. Interestingly, our data suggest that H2O2 treatment results in more defined transcript localization; a large cohort of transcripts that are classified as mixed in control, are localized after treatment. In other words, many of the transcripts that are roughly equally divided between the nucleus and cytoplasm under control conditions are either more stringently retained in the nucleus or more thoroughly exported to the cytoplasm after treatment. We find 1104 transcripts moving to nuclear localization and 1628 moving to cytoplasmic localization, evidence of an overall increase in the number of transcripts showing localization after exposure to oxidative stress (Figure 2D).

We also examined expression changes due to oxidative stress, through differential expression analysis using DESeq (Anders and Huber, 2012). We looked at each fraction separately, to identify expression changes specific to nuclear or cytoplasmic fractions after treatment, and we grouped differentially expressed transcripts by degree of change, so as to more thoroughly examine the effect of oxidative stress. We found that, while a subset of differentially expressed transcripts show a small degree of change after treatment, most transcripts with significant changes are either turned completely on or off (Figure 3). Our analysis of the nuclear fractions identified 611 coding transcripts significantly changed following H2O2 treatment. Of the 318 transcripts upregulated after treatment, 247 were not expressed in the untreated nuclear fractions, and of the 293 transcripts downregulated, 262 show no expression after treatment. We found slightly fewer transcripts with significantly altered expression after treatment in the cytoplasmic fractions, with a total of 559 coding transcripts showing changes. There were 277 transcripts upregulated after treatment, 202 of which were not expressed in the untreated control cytoplasmic fractions, and 282 transcripts downregulated, with 253 showing no expression after treatment. These data indicate that, regardless of the fraction analyzed and direction of change, the vast majority of transcripts with significantly altered expression are either turned on or off completely.

**Figure 3.**
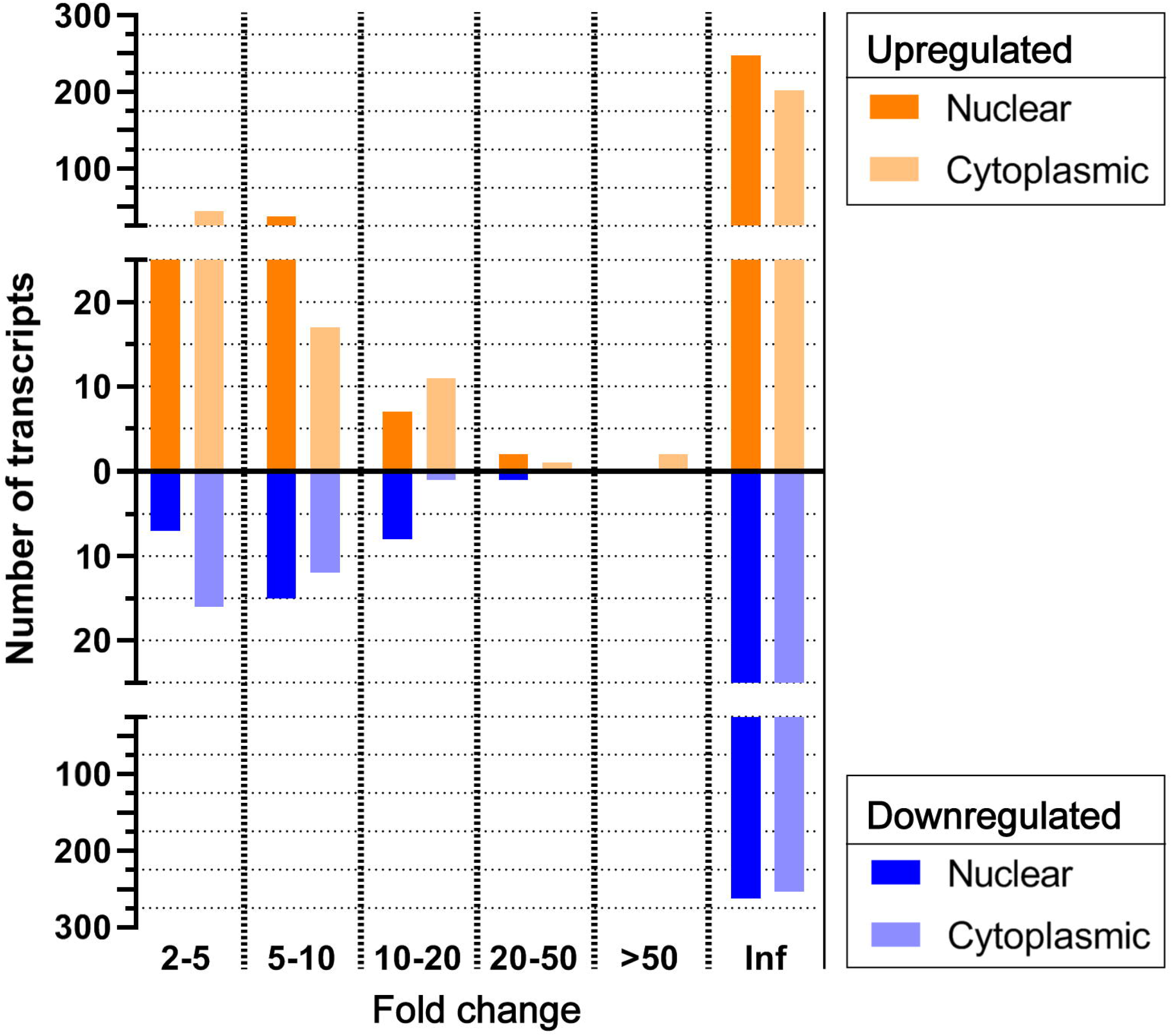
Majority of differentially expressed transcripts are turned on or off by peroxide treatment. Transcripts differentially expressed after peroxide treatment in both nuclear and cytoplasmic fractions. Transcripts are graphed by direction of change and binned by fold change. In both the nucleus and cytoplasm, most DE transcripts are either turned on (have no expression in control) or turned off (have no expression after treatment) as indicated by the infinite (Inf) fold change bin.

### 3. Specific Genes of Interest

Given the importance of the RPE to retinal function and disease pathogenesis, we specifically analyzed the localization of three sets of genes: those involved in AMD, those involved in IRDs, and the RPE markers. In all three of these gene sets, we see an increase in overall localization after treatment, with more noticeable changes in the cytoplasmic fraction (Figure 4). The 51 genes with potential involvement in AMD are composed of 853 transcripts. Of these, 51 transcripts are localized to the nucleus and 7 to the cytoplasm in control iPSC-RPE. Following treatment, there is a slight increase in the number of transcripts localized to the nucleus (68) and over twice as many localized to the cytoplasm (17). This trend is even more evident in the 2299 transcripts expressed from 227 IRD genes (Daiger et al., 2013). In these genes, we again see a notable increase in localization to the nucleus after treatment, with 61 nuclear localized transcripts in the control and 97 after treatment. Moreover, nearly 3 times as many transcripts are localized to the cytoplasm after exposure to H2O2; we found 13 in the control compared with 36 after treatment. The RPE signature gene list is partially composed of genes from the other two sets, and consists of 86 genes expressing 958 transcripts. Here, we see roughly equal numbers of transcripts localized to the nucleus (40 control and 41 treated), but a striking 10 times as many transcripts localized to the cytoplasm in the treated relative to the control (10 control and 97 treated).

**Figure 4.**
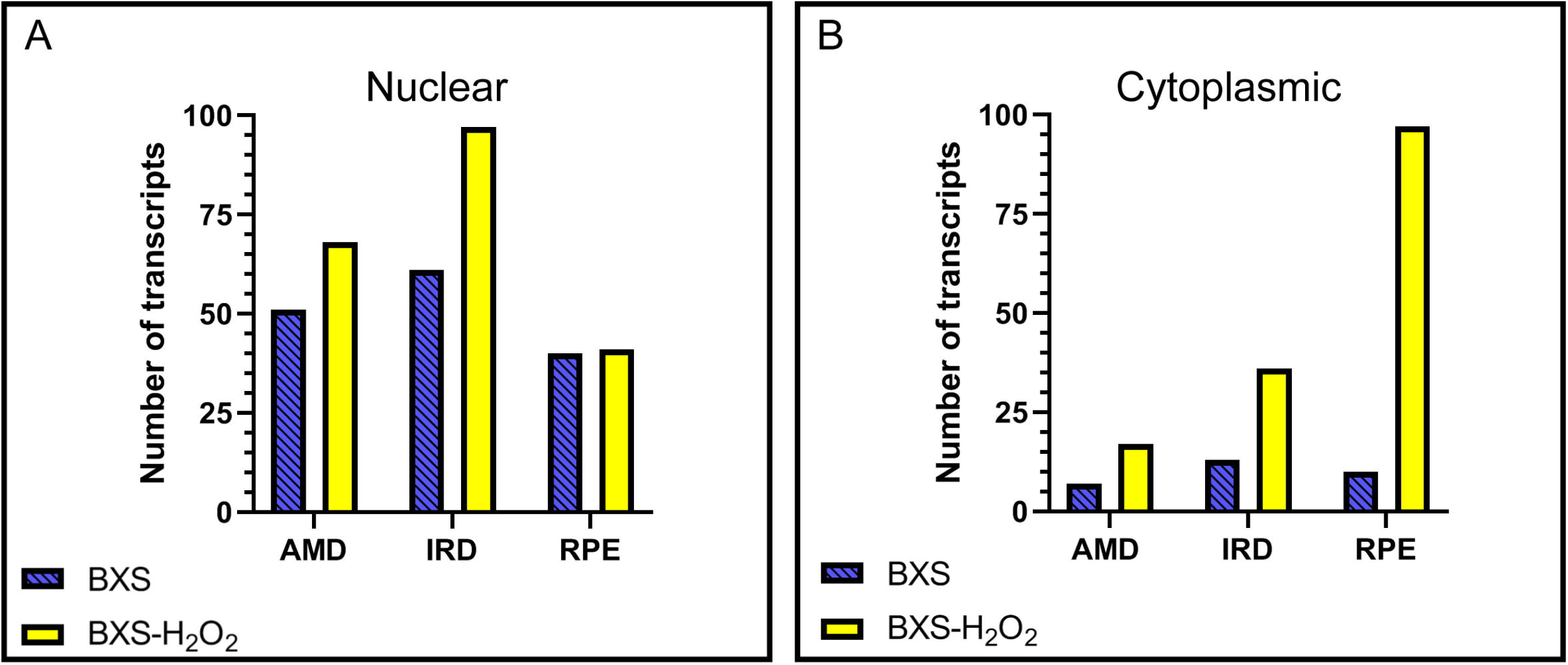
Localization of transcripts involved in AMD and IRDs, as well as RPE marker transcripts. Transcripts involved in AMD, IRDs, and RPE markers that localize to the nucleus (**A**) and cytoplasm (**B**) in control and treated samples.

Our in-depth analysis of transcripts from these gene lists revealed that the majority of transcripts localized to the nucleus in the control cells are also nuclear localized after treatment (94% AMD, 79% IRD, 78% RPE), while we see far fewer of the transcripts localized to the cytoplasm in the control continue to localize to the cytoplasm after treatment (14% AMD, 10% IRD, 38% RPE). The increase in cytoplasmic localized transcripts overall, coupled with the fact that few transcripts localized to the cytoplasm in control continue to be similarly localized after treatment warrants further exploration. As with the whole transcriptome, we find that a majority of the localization changes in these genes are a result of transcripts that are mixed in control cells moving to cytoplasmic localization after exposure to H2O2. Notably, we see multiple isoforms from each gene with altered localization. From the set of AMD genes, 12 transcripts from 5 genes move from mixed to cytoplasm after treatment, representing 10% of the genes in this list. Of the IRD genes, 23 transcripts from 11 genes shift localization from mixed to cytoplasm, representing 5% of this list. A full 20% of the RPE gene list shows a shift from mixed to cytoplasmic localization, with 89 transcripts from 18 genes moving in this way. This dramatic shift in transcripts localized to the cytoplasm after H2O2 treatment requires more in-depth analysis to determine the potential role these changes may play in the RPE and in AMD and IRD disease pathogenesis.

## Discussion

RNA retention in the nucleus is a mechanism for controlling protein abundance in individual cells (Bahar Halpern et al., 2015; Chin and Lécuyer, 2017; Kallehauge et al., 2012; Stoeger et al., 2016). This mechanism affords the cell the ability to quickly respond to changing environmental conditions, as has been shown for heat stress and hypoxia. Conversely, aberrant RNA retention can harm the cell, leading to disease pathogenesis, such as in myotonic dystrophy, Alzheimer’s Disease, and glial cell tumors (Comincini et al., 2006; Larkin and Fardaei, 2001; Mastroyiannopoulos et al., 2010; Sun et al., 2015). By evaluating the whole coding transcriptome, we found that oxidative stress causes changes in RNA localization, and these are especially evident in genes known to cause retinal degeneration.

Interestingly, our data show that mRNA export from the nucleus is more nuanced than the dogmatic view that mRNA is quickly processed and transported out of the nucleus following transcription. While it is known that some mRNAs undergo processing in addition to capping, splicing, and polyadenylation, and these extra processes will cause retention of the transcript, our data suggests that retention of mRNA is much more ubiquitous than previously understood (Chen and Carmichael, 2009; Prasanth et al., 2005). Specifically, our results contrast with the findings of Ulitsky, et al., (2015), where they found a majority of mRNA transcripts localized to the cytoplasm in the MIN6, mouse insulinoma pancreatic beta cell line, while only 30% of mRNAs were retained in the nucleus (Bahar Halpern et al., 2015). Further, in control mouse liver cells, they observed 13.1% of mRNAs to be retained in the nucleus. In contrast, we observed roughly 59% of localized transcripts retained in the nucleus in our control iPSC-RPE line, while H2O2 treatment led to 62% of localized mRNAs being retained in the nucleus. The differences observed between the two studies can be attributed to multiple factors. First, it is possible, and likely, that different cell types retain and/or export mRNA differently based on their needs. Second, species differences may play a role, as our study was performed in human cells, while the former was performed in mouse. Finally, cell origin could factor into the differences. The MIN6 cell line is derived from cancerous cells, while the liver cells were primary cultures. Reprogramming and differentiation, as was performed for the iPSC-RPE, may introduce variation, as well. With respect to gene expression, we have previously shown that while iPSC-RPE are mostly similar to native RPE, differences do exist (Au et al., 2017). These differences may carryover to mRNA retention and export.

The cellular consequences of RNA retention are situation dependent. For example, under normal environmental and cellular conditions, RNA retention can serve to buffer bursts of transcription as a mechanism to maintain proper protein levels (Bahar Halpern et al., 2015; Kallehauge et al., 2012; Stoeger et al., 2016; Wegener and Müller-McNicoll, 2018). Further, retaining excess transcripts in the nucleus can provide flexibility to the cell for rapidly responding to a stressor (Zander et al., 2016). On the other hand, abnormal retention, and conversely, abnormal transport to the cytoplasm, can lead to cytotoxicity (Redford-Badwal et al., 1996; Wegener and Müller-McNicoll, 2018). Given the high oxidative load experienced by the retina under normal conditions, we set out to understand the role oxidative stress has on localization of genes necessary for proper maintenance and function of the RPE. For example, we found that transcripts for both *IFT172* and *NPHP4* are not exported to the cytoplasm following H2O2 treatment. Both genes are involved in cilia formation, like those found in the apical processes of the RPE, and recessive mutations lead to retinitis pigmentosa, among other systemic pathologies. Similarly, *NPLOC4,* which is implicated in AMD risk, is also not exported to the cytoplasm following oxidative stress. On the other hand, export of an isoform of *RGR* is increased after treatment.

Disease pathogenesis is likely to be a complicated process, and not solely attributable to mutations in a single gene. Rather, it is highly probable that other factors, both internal and external, contribute to disease progression. Our study is the first to evaluate the localization of mRNA in the RPE under a ubiquitous environmental stressor. This work adds to the growing evidence that alters the paradigm of rapid mRNA export from the nucleus. Further, while localization changes of mRNA from disease-causing genes can potentially be implicated in disease progression under oxidative stress, it is also possible that these changes occur to protect the cell. Our data suggests a complicated, multifaceted impact of oxidative stress on altered nuclear retention, with promising potential for a role in retinal disease pathogenesis, and future studies are needed to fully explore these possibilities using disease models or primary tissue.

## Author Contributions

TJK and MHF designed the study. TJK and MHF performed experiments. EDA and MHF analyzed and interpreted the data. MHF, EDA, and TJK wrote and edited the manuscript.

## Data Availability Statement

The datasets generated for this study can be found in the GEO REPOSITORY, GEO accession GSE158909.

## Acknowledgments

Facilities and resources provided by the VA Western New York Healthcare System to MHF. MHF is also a Research Biologist at the VA Western New York Healthcare System, Buffalo, NY. Computational support was provided by the Center for Computational Research at the University at Buffalo. Next Generation Sequencing services were provided by the Genomics Core at the Children’s Hospital Los Angeles. The contents of this manuscript do not reflect those of the Department of Veterans Affairs or the U.S. Government.

## Contribution to the Field Statement

Regulation of RNA transcript distribution is an important contributor to cellular homeostasis. Although research in this field has seen great progress in recent years, it is still unclear the extent to which RNA localization is affected by cell type and extracellular stressors. Furthermore, it is unknown whether dysregulation of RNA localization could be involved in the pathogenesis of age-related macular degeneration (AMD) and other diseases of the eye. Here, to illuminate these issues, we explore the distribution of RNA in the retinal pigment epithelium (RPE), a tissue that is critical to the health and functioning of the eye. We demonstrate that, on the whole, RNA within normal RPE is more heavily localized to the nucleus than has been seen in other cell types and tissues. Additionally, we show that oxidative stress, which is a known risk factor for the progression of AMD, causes a more highly defined distribution of RNA within the RPE, with more transcripts localizing to either the nucleus or cytoplasm. Importantly, we also find that this shift in RNA localization extends to transcripts involved in eye disease and development, suggesting a connection between RNA localization and diseases of the eye.

