## Supplementary figures and images for "Oxidative stress alters transcript localization of disease-causing genes in the retinal pigment epithelium"

### Supplemental Figure 1

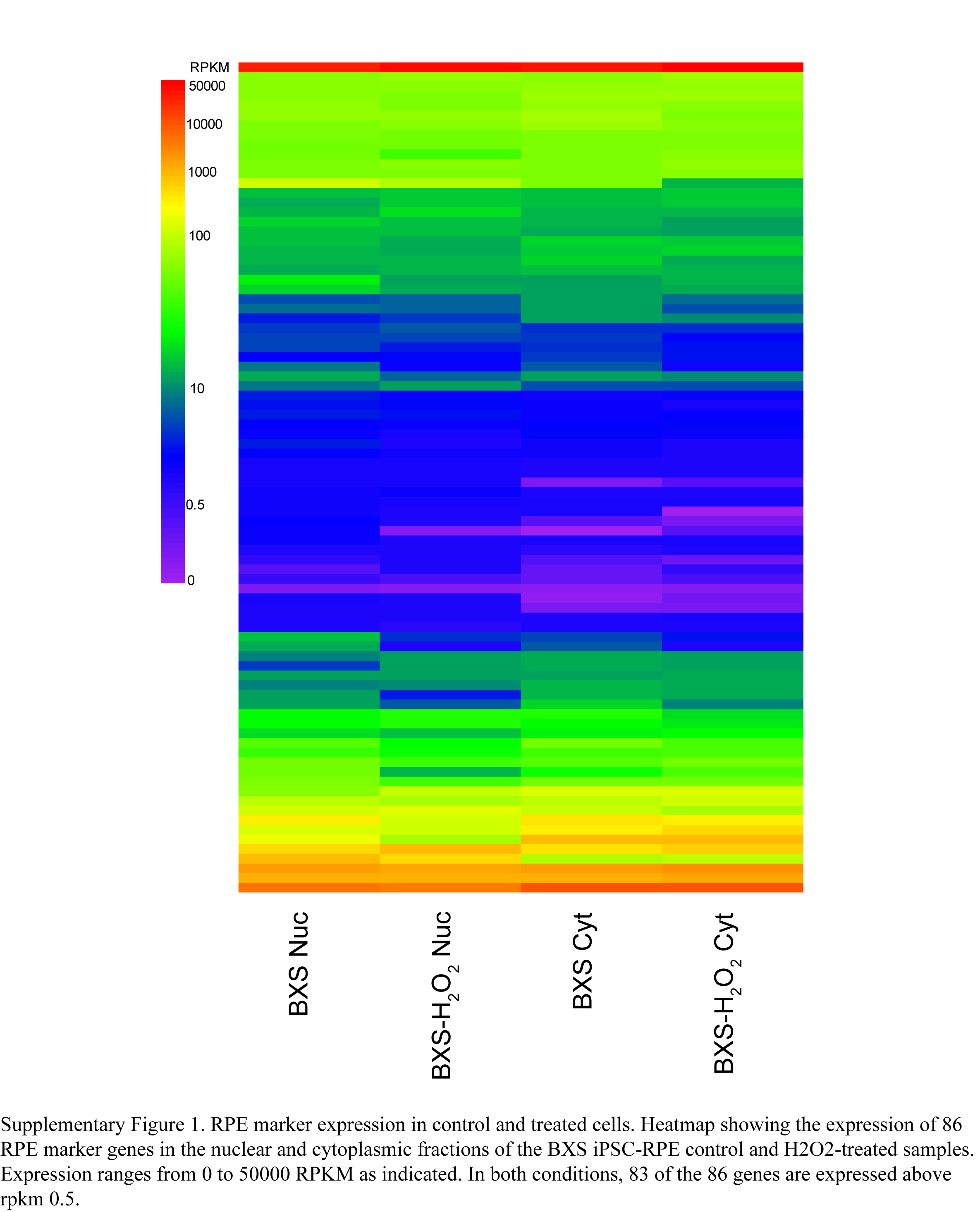
